# Circulating T-follicular helper and type I regulatory T cells have overlapping phenotypes in *P. falciparum* malaria and are maintained by parasite exposure

**DOI:** 10.64898/2026.06.24.734380

**Authors:** Mayimuna Nalubega, Damian A. Oyong, Dean W. Andrew, Megan S. F. Soon, Jessica R. Loughland, Zuleima Pava, Nicholas L. Dooley, Kenneth Musinguzi, Nankya Felistas, Isaac Ssewanyana, John Rek, Emmanuel Arinaitwe, Moses R. Kamya, Margaret E. Feeney, Prasanna Jagannathan, Christian R. Engwerda, Michelle J. Boyle

## Abstract

Immunity to *P. falciparum* malaria develops slowly, requiring repeated infection in areas of high transmission, and wanes rapidly in the absence of parasite exposure. Key to this immunity, is the development of antibodies which is supported by CD4⁺ T follicular helper (Tfh) cells that drive robust germinal centre responses. However, in malaria, the malaria-specific CD4⁺ T cell compartment in peripheral blood is dominated by Type 1 regulatory T cells (Tr1), which produce high levels of IL-10 in response to parasites. Tr1-like Tfh cells (Tfh10) have been reported in several settings of repeated antigen stimulation but have not been investigated in malaria. Here we used single-cell RNA sequencing and multiparameter flow cytometry to characterise malaria-specific Tfh and Tr1 cells in a longitudinal cohort of highly exposed individuals and assessed their persistence after transmission interruption. Malaria-specific Tfh and Tr1 cells shared overlapping profiles, and Tr1 cell-like transcriptional signatures and phenotypes were detectable within the Tfh cell compartment. Tfh10 cell subsets were the dominant phenotype of malaria-specific Tfh cells. Following disruption of malaria transmission, the frequencies of malaria-specific Tr1 and Tfh10 cells declined. These findings highlight a close relationship between Tfh and Tr1 cells and show that the Tfh cell compartment in malaria is dominated by Tfh10 cells. The rapid waning of these cells in the absence of continuous exposure is consistent with requirements of persistent antigen in maintaining regulatory CD4 T cell phenotypes.

## Introduction

Malaria, caused by *Plasmodium falciparum* infection, remains a significant health burden globally and control efforts have failed to reduce incidence since 2015 ^1^. In areas of high transmission, immunity to malaria develops slowly, and children experience repeated episodes of infection and disease before becoming clinically immune ^2^. Further, protective immunity is short-lived, and in the absence of continuous exposure previously immune individuals become susceptible to disease upon re-infection ^3,4^. CD4+ T cells are key mediators of immunity from malaria, with malaria specific CD4+ T cells having roles in parasite clearance, regulating inflammation and providing help to B cells ^5,6^. These functions are mediated by IFNγ producing Th1 cells, type 1 regulatory (Tr1) cells, and T-follicular helper (Tfh) cells respectively, however recent research has identified significant plasticity and overlapping phenotypes between these CD4+ T cell subsets ^7–12^.

In children with high exposure to malaria, Tr1 cells dominate the malaria specific compartment ^7,8,11–13^. Tr1 cells express IL-10, often together with IFNɣ, along with co-inhibitory receptors that mediated regulatory functions ^5^. Malaria specific Tr1 cells emerge early in malaria infection and may have roles in controlling immunopathogenesis, possibly at the expense of parasite clearance ^8^. Recently we have shown that malaria-specific Tr1 cells share transcriptional, phenotypic and clonal overlap with Tfh cells in controlled human malaria infection ^10^. Tfh cells are a specialised CD4+ T cell subset, expressing CXCR5 and PD1 which function within the germinal centre to support the development of humoral immunity ^17,18^. To drive the diversity of antibodies required for protection across pathogens, Tfh cells have diverse phenotypes, with specific subsets providing B cell help in a context specific manner ^19–22^. Human studies leveraging peripheral blood to analyse circulating Tfh cells that provide a snap shot of germinal centre Tfh from secondary lymphoid sites ^21,22^, have largely focused on analysing Tfh cell diversity based on expression of CXCR3 and CCR6 to identify Tfh1 (CXCR3+CCR6-), Tfh2 (CXCR3-CCR6-) and Tfh17 (CXCR3-CCR6+) cells ^18^. We have shown that in malaria both Tfh2 cells and a subset of Tfh1 cells that lack cytotoxic markers and express CCR7, are associated with antibody development in controlled human malaria infection models ^9,23^. Further, in an area of intense perennial transmission, both activated Tfh1 and Tfh2 cells are increased in children with the highest levels of functional antibodies ^24^. However, consistent with the overlap of Tr1 and Tfh cells in controlled human malaria infection ^10^, Tfh cells can also express IL-10, and Tr1-like Tfh (Tfh10) cells have been reported in multiple non-malaria experimental models and human studies ^24–30^. The Tfh10 cell subsets have yet to be investigated in malaria infection, but given the dominance of malaria-specific Tr1 cells, this knowledge gap needs to be addressed.

Here we investigated Tfh and Tr1 cells and the overlap of these subsets in individuals from an area of East Africa with intense year-round malaria exposure with scRNA/TCRseq and high dimensional flow cytometry, identifying malaria-specific CD4 T cells with activation induced markers (AIM) assays. We show significant transcriptional and phenotypic overlap between Tr1 and Tfh cells and show that Tfh10 cells dominate the Tfh cell response in malaria. In the absences of continuous malaria exposure, both Tr1 and Tfh10 cells decline, suggesting these phenotypes are driven by high and continued antigen exposure, and consistence with roles of these cells in maintenance of tolerogenic immunity^8^.

## Results

### Malaria-specific Tfh and Tr1 cells have overlapping transcriptional signatures

To investigate malaria-specific Tfh and Tr1 cells, we leveraged samples from children in a cohort in Uganda with intense year-round malaria exposure. We first analysed the entire CD4 T cell compartment in three children who had samples available at four time points over a period of 9-14 months, along with samples from 5 adults taken from a single time point. PBMCs were stimulated with parasite infected RBCs to identify antigen specific cells based on activation induced markers, CD69 and OX40 ^9,32,33^ (AIM+ cells). All AIM+ cells were sorted and analysed by 10X 5’ scRNA/TCRseq, along with a proportion of AIM- cells from each individual and timepoint (Figure 1A/B, Supplementary Figure 1A/B). Following integration by sequencing batch, donor and antigen specificity, 11 CD4 T cell subsets were identified and annotated based on differential gene expression (DEGs) (Figure 1B, Supplementary Figure 2A/B, Supplementary Table 1). This approach identified two populations of Tfh cells (annotated resting Tfh and Tfh cells) which expressed *CXCR5*, *IL21*, *ICOS* and *PDCD1* (encoding PD-1) (Figure 1C). Resting Tfh cells also expressed markers of central memory cells (*CCR7* and *CD27*), consistent with the transcriptional profile of these cells in recent study in children from another Ugandan cohort ^8^. Tr1 cells were annotated based on high expression of *CTLA4, MAF*, and *LAG3.* These Tr1 cell-associated marker genes were also expressed at lower levels within the Tfh cell cluster. Two populations of cells were annotated as Th1 and Cytotoxic-Th1 cells, based on expression of *CXCR3.* The Cytotoxic-Th1 cell cluster also expressed high levels of *GZMB, GZMA, NKG7,* and *GNLY.* Consistent with our recent findings ^10^, there was significant overlap between Tfh and Tr1 transcriptional profiles, with 317 genes overlapping between the two subsets (Figure 1D). However, key markers of Tfh cells *CXCR5, IL21,* and *CD40LG* where all higher in the Tfh cluster, while the Tr1 cluster expressed higher levels of *IL10, HAVCR2, PRDMI* (encoding BLIMP1), *MAF, TMEM173* (encoding STING, a key driver of Tr1 cells ^34^), *LAG3* and *CTLA4* (Figure 1E). Further, analysis comparing the Tfh and Tr1 cell clusters to each other identified 1175 DEGs (Supplementary Table 2). DEGs between Tfh and Tr1 cells included relatively upregulated expression of *CXCR6, GZMA, GZMK* in Tr1 cells, all of which were identified as Tr1 cell marker genes in a recent analysis in children from the same study area ^7,8^ (Supplementary Table 2). Other annotated clusters included Th2 cells expressing *GATA3, CCR4* and *IL4R*, and a Th2-IFN cell cluster which expressed Th2 genes and interferon signalling genes *(IFI44L, ISG15, IF16, MX1, XAF1)*; Th17 cells expressing *CCR6,* and *RORA*; Treg cells which expressed *FOXP3, IKZF2,* and *TIGIT;* and two subsets of central memory (CM) cells, one of which also expressed IFN signalling genes and was annotated CM-IFNs (Figure 1C).

**Figure 1:**
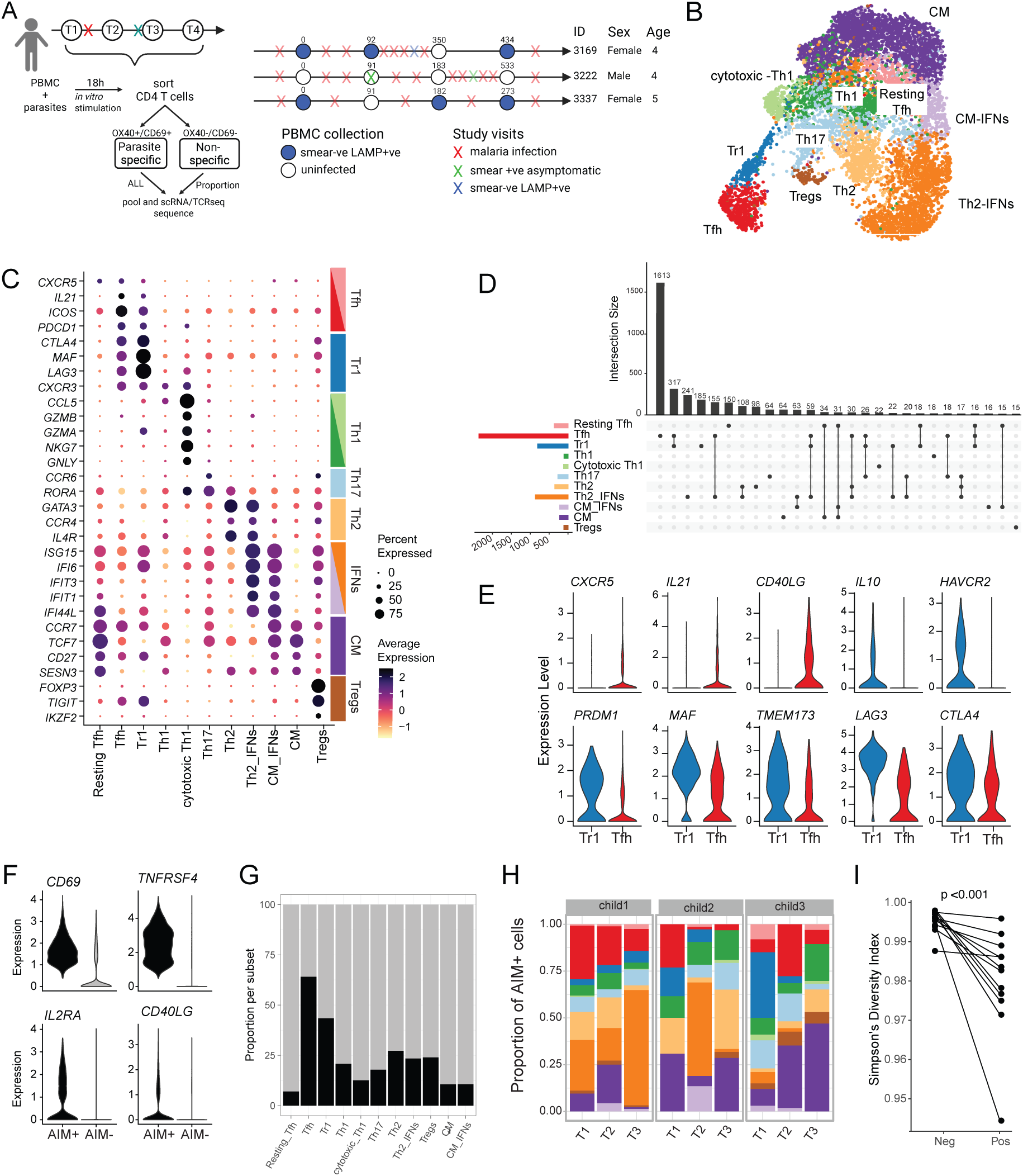
scRNAseq of malaria-specific CD4 T cells. **A)** PBMCs were stimulated with parasite infected RBCs for 18 hours to identify malaria specific cells with Activation Induced Marker assays. Malaria-specific cells were enriched by cell sorting OX40+CD69+ CD4 T cells. All malaria specific cells were sequenced, along with a proportion of AIM- cells. PBMCs from 3 children in an area of intense malaria transmission were analysed. PBMCs were collected when children were uninfected, or asymptomatically infected (detected by blood smear or LAMP), with at least one clinical malaria episode between sampling timepoints. **(B)** UMAP visualisation of CD4 T cell clusters. **C)** Dot plot of average expression of DEGs used for cluster annotation. **D)** Upset plot of the number of shared and specific marker genes across CD4 T cell clusters. The bar graph on the right shows the total number of DEG used to annotated each subset. **E)** Violin plots of key marker genes for Tfh and Tr1 cells in both subsets. **F)** Violin plots of TCR engagement genes in AIM+ (malaria specific) and AIM- cells. **G)** Proportion of AIM+ malaria-specific cells within each cluster. **H)** Proportion of each CD4 T cell subset in AIM+ cells in each child across timepoint. **I)** Simpson diversity index of AIM+ compared to AIM-cells. See also Supplementary Figure 1 and 2, and Supplementary Tables 1 and 2.

To analyse malaria-specific CD4 T cells within the dataset, we identified TCR engaged CD4 T cells based on expression of activation induced markers *CD69* and *TNFRSF4* (encoding OX40), which identified 2968 malaria - specific AIM+ CD4 T cells (35.6% of sequenced cells) (Figure 1F). These malaria-specific AIM+ cells also expressed elevated expression of *IL2RA* and *CD40LG* (encoding CD25 and CD40L, respectively), consistent with TCR engagement (Figure 1F). Malaria-specific AIM+ CD4 T cells were identified across the CD4 T cell compartment and were relatively enriched in Tfh and Tr1 cells (Figure 1G, Supplementary Figure 2D). At the individual level, malaria specific cells were phenotypically diverse, however Tfh and Tr1 cells comprised a large proportion of antigen specific cells (Fig. 1H). Within each child, the proportion of each cluster varied over time. However, there were no apparent consistent changes, possibly due to the relatively short time frame of sampling (Figure 1H). Across CD4 T cells, TCR sequences with complete TRA/TRB genes were captured for 5020 (60.3%) cells. Despite our approach to enrich parasites specific cells using the AIM, the majority of cells were single TCR clones (92.5%), and clonal expansion was only detected in 484 cells across 176 unique clones ranging from 2-35 cells per clone. Nevertheless, consistent with a higher proportion of expanded clones in the parasites specific AIM+ cells compared to non-antigen specific cells, Simpson’s diversity index was significantly lower in the malaria specific compartment (Figure 1I). While the detection of clonally expanded cells was low, of the 4 clones found within the Tfh cell cluster, 3 of them shared clonal overlap with Tr1 cells.

### Malaria specific Tfh cells are dominated by Tr1-like transcriptional signatures

To further examine the overlap in transcriptional profiles between malaria-specific Tr1 and Tfh cells, we next took a complementary approach, sequencing Tfh cells sorted from CD4 T cells based on CXCR5 expression following parasite stimulation (Figure 2A). Tfh cells were analysed from 5 different children from the same study site at four timepoints over 9-18 months, and 3 adults at a single time point (Figure 2A, Supplementary Figure 3). Within sorted and sequenced Tfh cells, 7 clusters were identified following integration by donor (Figure 2B, Supplementary Figure 4A-D, Supplementary Table 3). A Tr1-like cluster, annotated Tfh10, was identified with high expression of Tr1 cell-associated genes including *LAG3*, *MAF, CTLA4,* and *TIGIT* (Figure 2C). These cells also had high expression of Th1 cell-associated genes *CXCR3,* and cytotoxic genes *CCL5, GZMK*, *GZMM,* and *CTS7.* Additionally, Tfh10 had reduced expression of memory-associated genes *CCR7* and *SELL,* suggesting an effector memory phenotype. Other Tfh cell clusters were annotated as Tfh2 (expressing *GATA3* and *IL4R),* an IFN1 cluster (enriched in Type I IFN genes), while remaining clusters had unclear classifications (Figure 2B/C).

**Figure 2:**
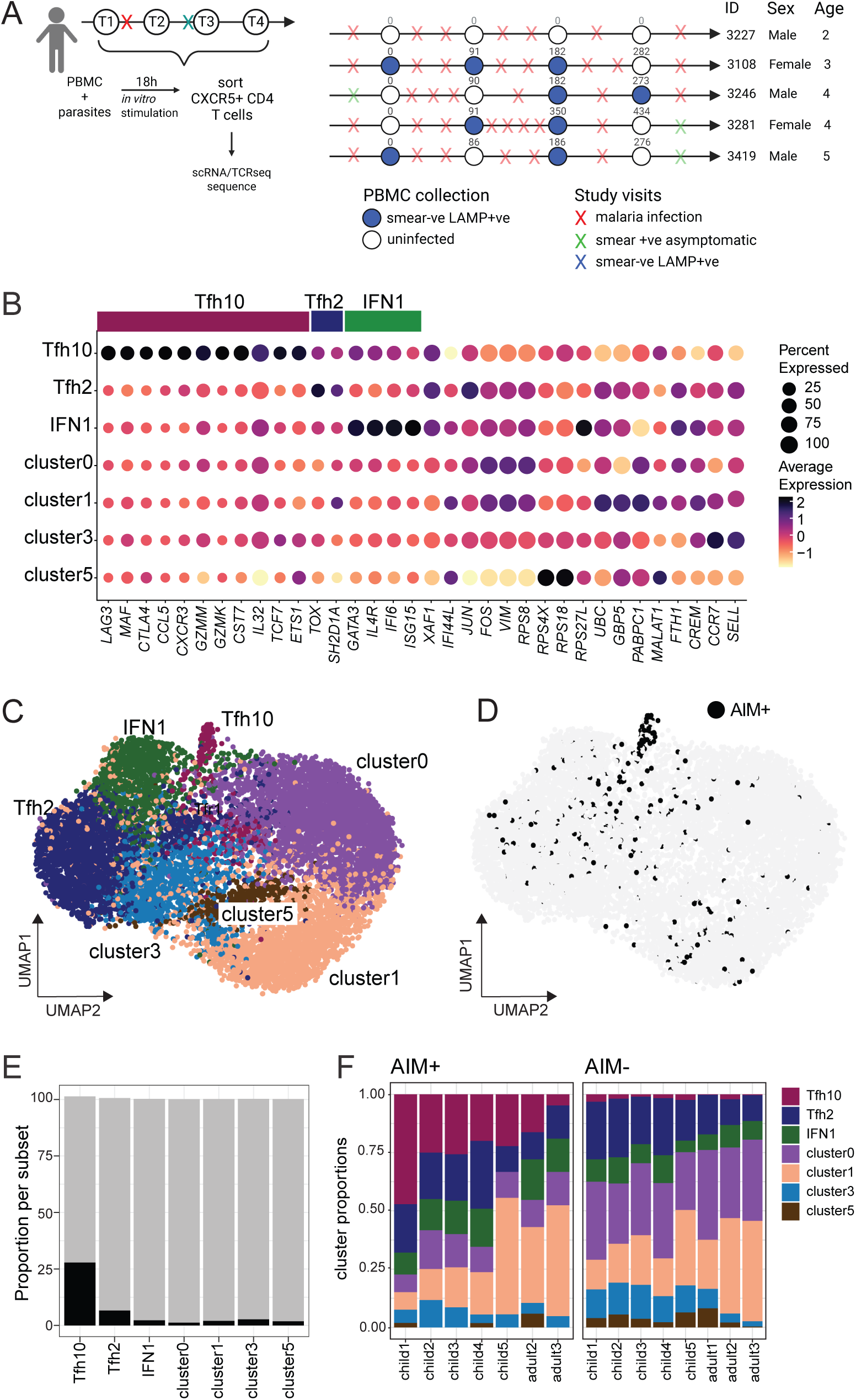
Malaria specific Tfh cells are dominated by Tfh10 phenotypes. **A)** PBMCs were stimulated with parasite infected RBCs for 18 hours and Tfh cells (CXCR5+ CD4 T cells) were sorted and sequenced. PBMCs from 5 children in an area of intense malaria transmission were analysed, collected when children were uninfected, or asymptomatically infected (detected by blood smear or LAMP), with at least one clinical malaria episode between sampling timepoints**. B)** UMAP visualisation of Tfh cells from 10663 cells. **C)** Dot plot of the average expression of curated marker genes for each Tfh cell cluster. **(D)** UMAP visualisation of malaria-specific AIM+ Tfh cells (black) within data set. **(E)** Proportions of malaria-specific cells within each Tfh subsets. **F)** Proportion of each subset within malaria-specific AIM+ and non-specific AIM- cells within each donor. Adult1 did not have any AIM+ cells within data set. See also Supplementary Figure 4, 5 and Supplementary Tables 3.

To identified malaria-specific cells with the Tfh data set, we examined expression of TCR engagement genes and annotated cells as malaria-specific based on expression of *CD69* with either *TNFRSF4, IL2RA or TNFRSF9* (encoding OX40, CD25 and CD137, respectively) (Supplementary Fig. S4E). This approach identified, 327 cells (3%) as malaria specific AIM+ Tfh cells. Malaria-specific cells were significantly enriched within the Tfh10 cell cluster, made up to 50% of all malaria-specific cells and were almost absent in the non-antigen-specific compartment (Figure 2D/E/F). Amongst the data, TCR sequences with complete TRA/TRB genes were captured for 6405 (59.7%) cells. However, detected clonal expansion was limited, with 99.2% and 100% of cells present as singletons in the AIM- Tfh and AIM+ Tfh compartment, respectively. Thus, analysis of clonal sharing between Tfh subsets was not possible. However, together data identify a strong Tr1-transcriptional signature within Tfh cells in this population of highly exposed children, and that malaria-specific Tfh cells are enriched in Tfh10 cells.

### Tfh and Tr1 cell expansion is malaria specific

Transcriptional analysis of malaria-specific cells suggests significant overlap between Tr1 and Tfh, with Tfh10 dominating the Tfh cell compartment in malaria. To examine if this was malaria specific, or also occurred in other chronic infections, we next examined responses in additional children and adults from the same study area (children n=12, age median 4.58 years range 0.5-8.5, 66% female, and adults n=7 age median 33.2 years range 25.5-38.2, 100% female). To compare malaria response to other chronic pathogens, antigen specific cells were identified via AIM assays following stimulation with parasites and cytomegalovirus (CMV) protein (Figure 3A, Supplementary Figure 5A). CMV, which establishes a life time infection, infects nearly 100% of children in this study cohort ^33^. The frequency of malaria-specific CD4 T cells was higher than CMV-specific CD4 T cells and malaria-specific cells were higher in adults, but not different with age for CMV (Figure 3B/C).

**Figure 3:**
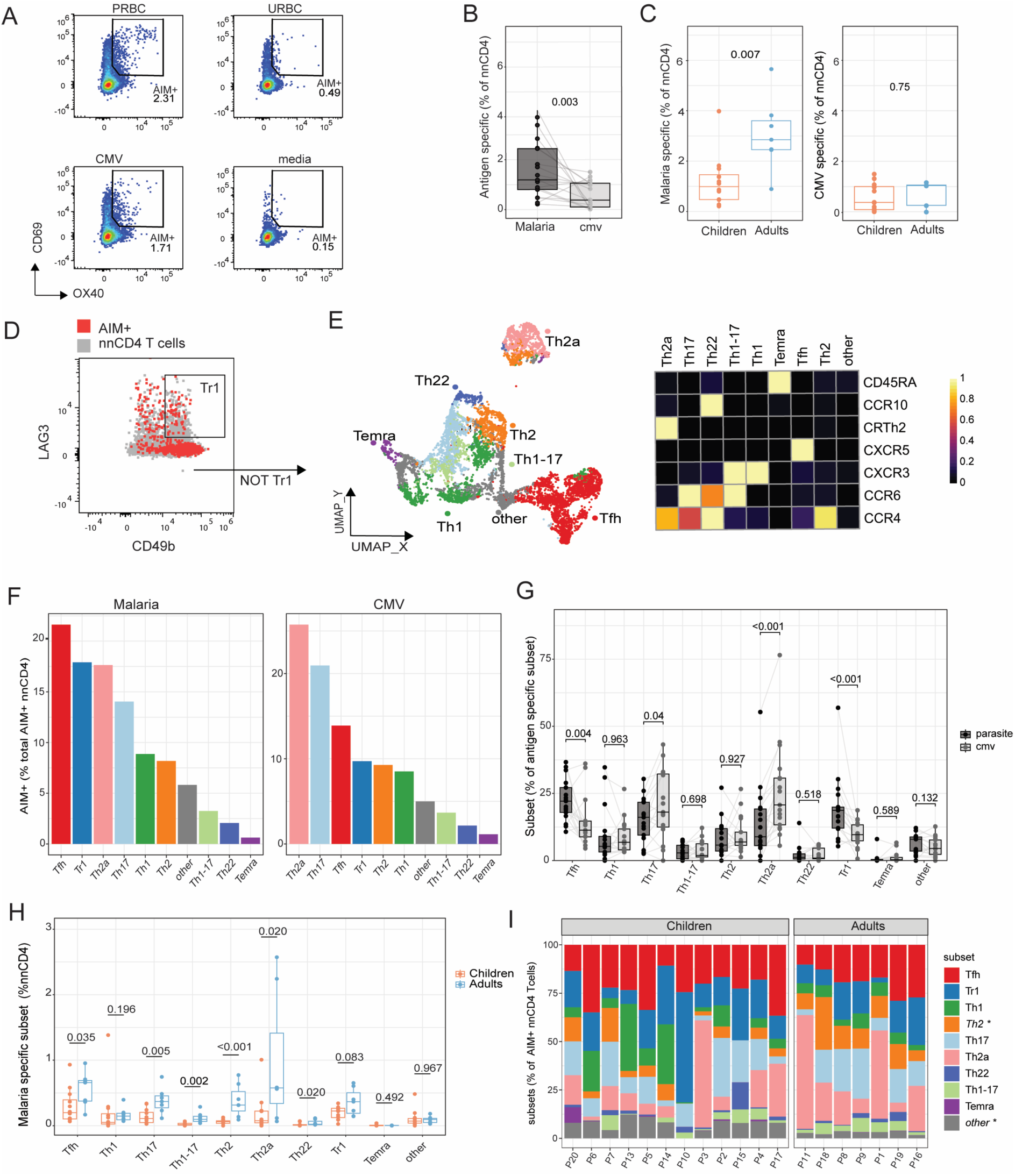
Comparison of malaria and CMV specific CD4 T cells. **A)** Representative gating of activation Induced Marker assays to identify malaria and CMV specific CD4 T cells following stimulation with parasite infected RBCs and CMV peptides. Responses from children (n=12) and adults (n=7) were analysed. (**B**) Frequency of malaria-and CMV-specific CD4 T cells (identified based on AIM expression) in non-naïve CD4 T cells (nnCD4) (**C**) Frequencies of malaria and CMV – specific CD4 T cells in children and adults. (**D**) Gating figure of Tr1 cells in nnCD4 and antigen-specific CD4 T cells identified by LAG3 and CD49b expression. (**E)** UMAP of CD4 T cell subsets identified by unsupervised clustering. Heatmap showing median expression of the various cell surface markers used in cell annotation. (**F**) Proportion of each subset amongst all antigen specific cells for malaria-specific and CMV-specific (**G)** Subsets as proportions of total malaria- and CMV- specific nnCD4 T cells. (**H**) Frequencies of each malaria-specific CD4 T cell subset in nnCD4 T cells in children and adults. (**I)** Proportion of each subset in malaria-specific response at individual level. Th2 and other cells were significantly different between children and adults. Box and whisker plots indicate first and third quartiles for hinges, median line and lowest and highest values no further than 1.5 IQR from hinges. P values are for comparisons between antigens are tested using Wilcoxon rank-sum test, and between children and adults using Mann-Whitney U test. See also Supplementary Figure 4.

CD4 T cell phenotypes were investigated amongst malaria and CMV antigen specific and non-specific cells based on expression of CXCR5, CXCR3, CCR6, CCR4, CRTh2, CCR10, CD45RA and CCR7, along with LAG3 and CD49b ^35^. Tr1 cell subsets could not be identified when clustering CD4 T cells based on expression of all markers, possibly indicating diverse co-expression of multiple chemokine receptors. As such, Tr1 cells were identified with manual gating as LAG3+CD49b+ cells, and remaining non-Tr1 cells clustered (Figure 3D). Amongst non-Tr1 cells, clustering identified Tfh (CXCR5+), Th1 (CXCR3+ CCR6-), Th2 (CXCR3-CCR6-CCR4+), Th17 (CXCR3-CCR6+), Th1-17 (CXCR3+CCR6+), Th2a (CRTh2+, CCR4+), Th22 (CCR10+ CCR4+ CCR6+), terminally differentiated effector memory CD4 T cells, TEMRA (CD45RA+), and a cluster which did not express any chemokine receptors annotated as ‘other’ (Figure 3E, Supplementary Figure 5C). Within each subset, the frequency of CD69+ OX40+ AIM+ malaria-specific or CMV-specific cells was quantified. The predominant subset of malaria-specific CD4 T cells was Tfh, followed by Tr1 cells, while for CMV the dominant responses were Th17 and Th2a (Figure 3F). Th17 cells were also detected within the malaria-specific CD4 T cell compartment, however it is likely these cells are a population of cytokine activated cells rather than antigen-specific ^36^ because Th17 CD4 T cells are also detected in malaria naïve individuals with AIM assays ^9^. Amongst the antigen-specific cells, the proportion Tfh and Tr1 cells was significantly higher for malaria, and Th17 and Th2a cells were significantly higher for CMV (Figure 3G). Comparing responses in children to adults, there was an age associated expansion of malaria-specific CD4 T cells, with adults having significantly higher frequencies of multiple subsets (Figure 3H). However, when considering the proportion of subsets within the malaria-specific response, few age-dependent differences were detected (Figure 3I, Supplementary Figure 4D). Together, these data show that the dominance of Tfh and Tr1 cells in malaria is likely pathogen specific, and that the entire malaria-specific CD4 T cell compartment expands with age in an area of high malaria transmission.

### Tr1 and Tfh cells phenotypes overlap in malaria specific CD4 T cells

In the above analysis, Tr1 cells were considered a distinct subset, gated based on LAG3 and CD49b prior to cell clustering. However, scRNAseq of CD4 T cells and Tfh suggested overlap between Tfh and Tr1 cells, and a dominant Tfh10 subset within the Tfh compartment. To assess this overlap phenotypically, we clustered Tr1 cells (identified as LAG3+/ CD49b+ with manual gating), based on expression of CXCR5, CXCR3, CCR6 and CCR4. Within the Tr1 cells, five clusters where identified, including Tfh cells (CXCR5), along with Th1 (CXCR3+CCR6-), Th2 (CXCR3-CCR6-CCR4+), Th17 (CXCR3-CCR6+) and Th17-17 (CXCR3+CCR6+) (Figure 4A, Supplementary Figure 6A/B). The proportions of Tr1 sub-clusters varied across individuals, however the majority of malaria-specific Tr1 cells had Tfh or Th1 cell phenotypes (Figure 4B, Supplementary Figure 6C).

**Figure 4:**
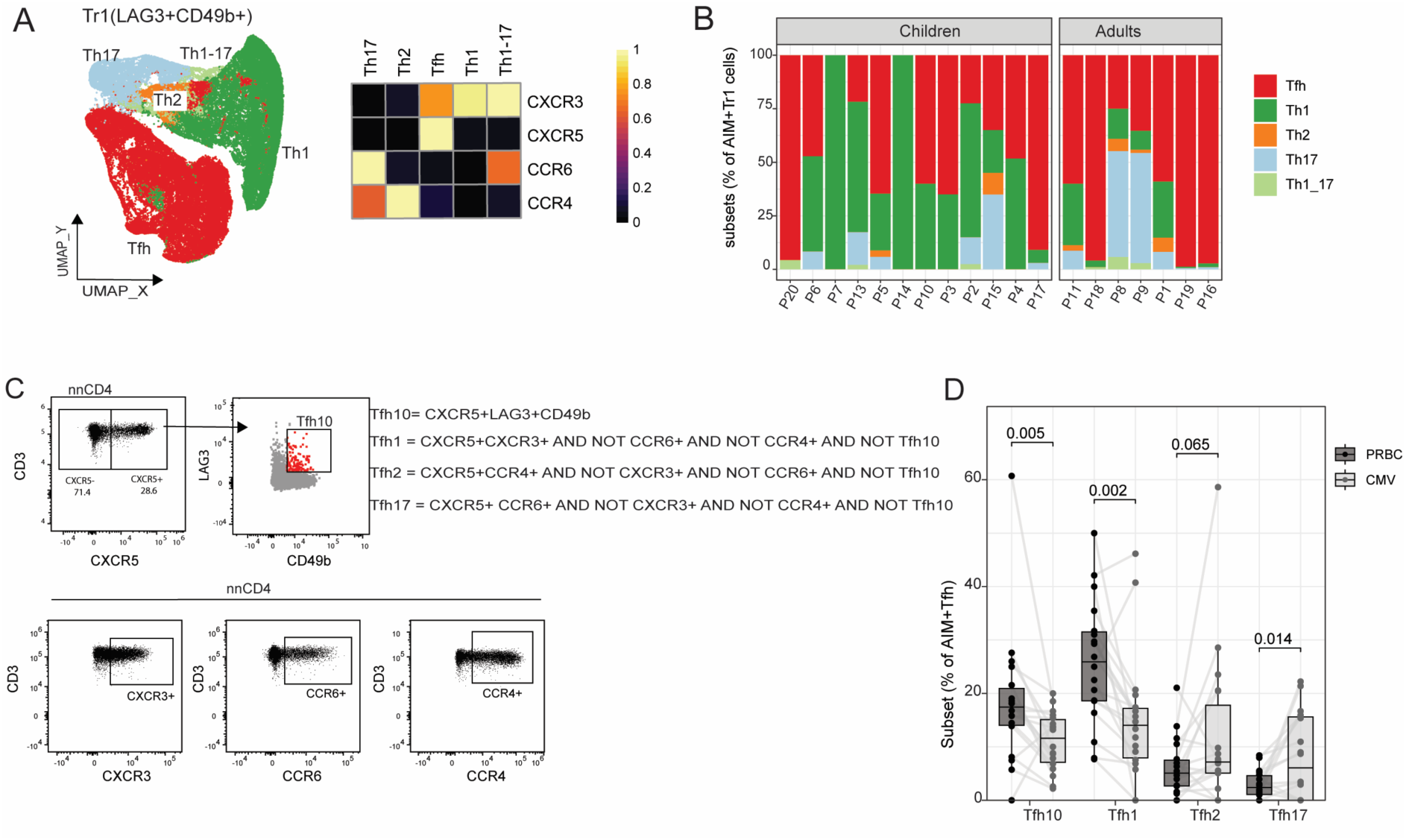
Regulatory Tfh cells in malaria-exposed children and adults. **A)** AIM+ and AIM- Tr1 CD4 T cells (LAG3+CD49b+) were clustered based on expression of CXCR3, CXCR5, CCR6 and CCR4. UMAP and heatmap of chemokine expression on identified clusters shown. **B)** Proportion of each subset amongst malaria-specific Tr1 cells across individuals. **C)** Gating strategy to identify Tfh, Tfh10 cells and other Tfh subsets by manual gating. **D)** Proportion of antigen-specific Tfh10 within all antigen specific Tfh cells in malaria and CMV. Box and whisker plots indicate first and third quartiles for hinges, median line and lowest and highest values no further than 1.5 IQR from hinges. P values are for comparisons between antigens are tested using Wilcoxon rank-sum test. See also Supplementary Figure 6.

Additionally, in a complementary analysis strategy, we re-analysed Tfh cells, identifying the entire Tfh cell compartment with manual gating as CXCR5+ CD4 T cells, and then identifying Tfh10 cells as LAG3+CD49b+ Tfh cells (Figure 4C). Within malaria-specific cells, Tfh10 cells were approximately 20% of all antigen-specific Tfh, consistent with frequencies detected within scRNAseq analysis (Figure 2E, 4D). The proportion of antigen-specific Tfh10 was significantly higher for malaria- compared to CMV-specific cells, suggesting that the expansion of Tfh10 is pathogen specific (Figure 4D). Tfh1 cells were the other dominant Tfh response in malaria (Figure 4D). Together, these data confirm the overlap of Tfh and Tr1 cells, and the malaria dependent expansion of Tfh10 cells within the Tfh compartment.

### Malaria exposure is required for the maintenance of malaria-specific Tr1 and Tfh10, independently of age

Finally, to assess if parasite exposure is required for maintenance of Tr1 and Tfh10 cells, we assessed malaria-specific CD4 T cells phenotypically within the same children (n=12) and adults (n=7) across three timepoints following the introduction of indoor residue spraying (IRS), which dramatically reduced transmission and infection incidence (Figure 5A). Focusing on malaria-specific Tr1, Tfh and Th1 cells, while large heterogeneity in frequencies across time was seen, there was clear decline in the frequencies of malaria-specific Tr1, Tfh and Th1 cells following IRS (Figure 5B). We modelled the decline in malaria-specific CD4 T cell subsets using a Tweedie Mixed effects Model to predict the impact of time (weeks since earliest assessed timepoint) during transmission interruption. Within the model, age was included as an interaction term, due to the reported impact of age on maintenance of immunity, including within this study site ^4,39^. We centred the model at age 10, an age where children in areas of high malaria transmission have developed immunity that protects from clinical malaria ^40^. Modelling showed that malaria-specific Tr1, Tfh and Th1 cells significantly declined over the period of IRS (Figure 5C/D, Supplementary Table 4). For Tr1 cells, the decline in malaria-specific Tr1 cells was only dependent on time, with no significant impact of age, suggesting Tr1 cells require constant antigen exposure for maintenance (Figure 5B/C). The decline of these cells is also consistent with a role in protective tolerogenic immunity, which is rapidly reduced in the absences of exposure ^3,4^. In contrast, for Tfh and Th1 cells, decline was significantly modified with age, with evidence for more rapid declines for children aged <10, and slower declines in individuals >10 (Figure 5B/D, Supplementary Table S4). The age dependent maintenance of Th1 and Tfh is consistent with patterns of antibody maintenance in this same study setting, where levels of antibodies maintained post IRS are higher in adults compared to children ^39^, suggesting a higher memory maintenance for these subsets with age. Declining frequencies of malaria specific cells were not seen for all subsets, for example, Th17 cells were not impacted consistent with these responses being non-specific bystanders ^37^. Additionally, there were slight increases in malaria-specific Th2 cell frequencies, possibly indicating that these responses are not all antigen specific, or alternatively, phenotypic plasticity between memory subsets.

**Fig 5:**
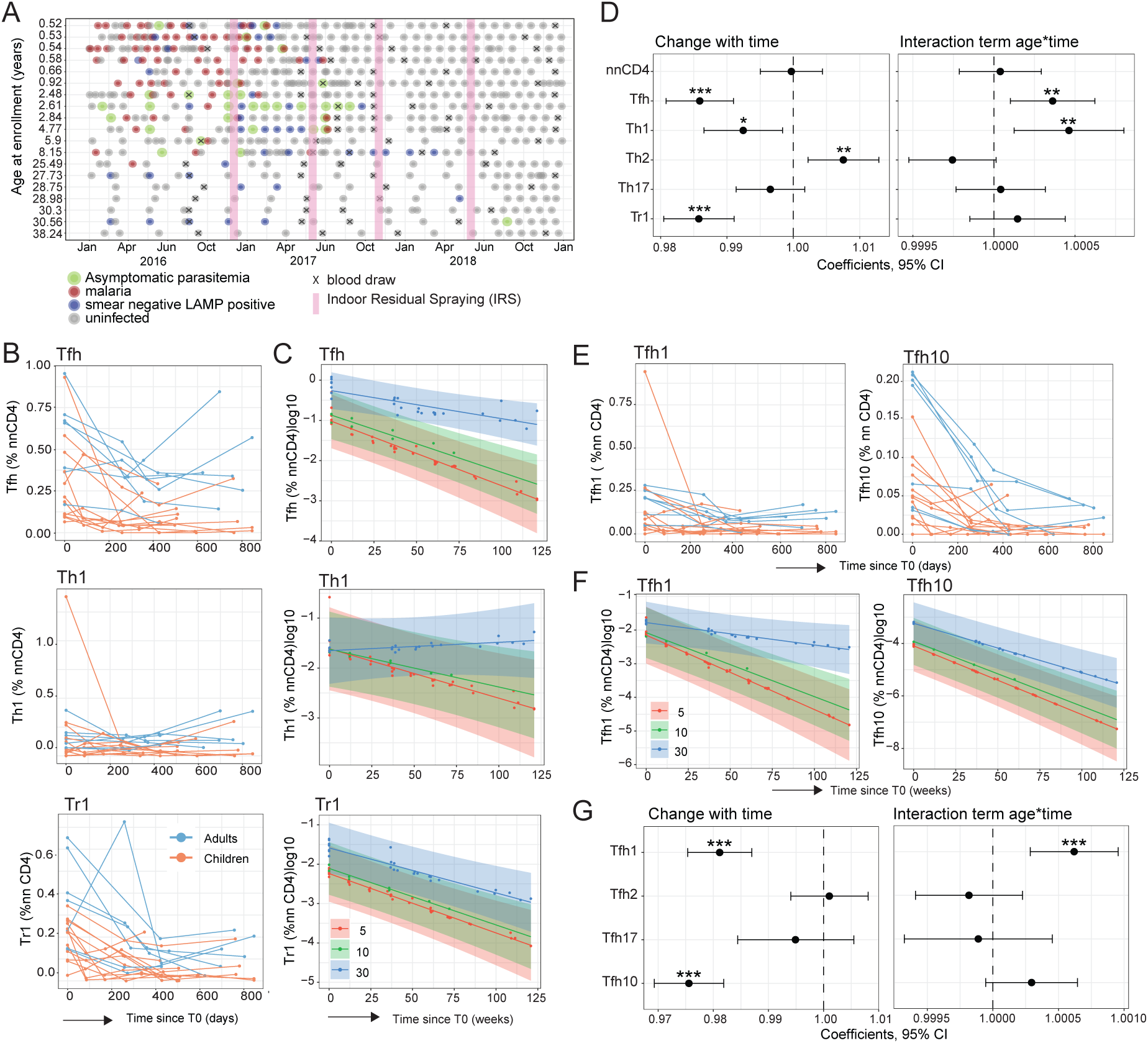
Effects of age and transmission disruption on malaria specific CD4 T cell subsets. (**A**) Timepoints of sample collection, IRS, malaria (symptomatic, asymptomatic and low-density infection [blood smear negative and LAMP positive]) amongst cohort. **A/E)** Longitudinal frequency of malaria-specific CD4 T cells (**A)** and Tfh cell (**E)** subsets as proportions of non-naïve (nn)CD4 T cells in children(red) and adults(blue). Data from the same individual are connected with a line. **B/F)** Tweedie model plots displaying the predicted log (frequency) of malaria-specific CD4 T cell subsets over time, conditional on fixed ages 5,10 and 30(age centred at −5,10,20), with 95% confidence bands. The dots are partial residuals from the model. The dots and lines are coloured by age. **C/G)** Effects of time during transmission disruption (Time since T0, weeks) and **(D/H)** the interaction of age (years) and time on malaria specific CD4 T cell subsets. Each point represents the estimated slope of cell log(frequency) change for age centred at 10 years, with horizontal lines indicating the 95% confidence intervals. The vertical dashed line represents no change in cell frequency. p is * < 0.05, ** < 0.01 and *** < 0.001

To assess if the decline of Tfh10 cells within the Tfh compartment was similar to the total Tr1 population, we analysed the frequencies of malaria-specific Tfh subsets identified by manual gating (within the CXCR5 compartment, Tfh10 LAG3+CD49b+ cells, and non Tfh10 classified as Tfh1 (CXCR3+CCR6-CCR4-), Tfh2 (CXCR3-CCR6-CCR4+), and Tfh17 (CXCR3-CCR6+CCR4-) (Figure 5E). Both malaria-specific Tfh10 and Tfh1 declined following IRS (Figure 5F). Additionally, while the observed decline in Tfh10 was age independent, the decline in Tfh1 was modulated by age. Taken together data highlight that the Tr1 and Tfh compartment dominates the malaria specific response in high endemic areas, and that maintenance of Tr1 cells, including Tfh10 within the Tfh compartment requires persistent parasite exposure.

## Discussion

CD4 T cells play a central role in malaria immunity, and Tr1 cells dominate the response in highly exposed children ^7,8,12,41–43^. Here, using scRNAseq and multiparametric flowcytometry analysis, we show that signatures of Tr1 cells are also dominant within the Tfh cell compartment, and the majority of malaria-specific Tfh cells having Tfh10 phenotypes. The dominance of malaria-specific Tr1 and Tfh10 responses was not a general characteristic of chronic infection and was significantly greater in malaria compared to CMV. Malaria-specific Tr1 and Tfh10 cells declined following reduced transmission, suggesting that these cells are maintained by antigen stimulation. Given the importance of both Tr1 cells in controlling symptomatic malaria, and Tfh cells in driving antibodies that suppress parasite growth, these findings have important implications for our understanding of the development and maintenance of clinical protection.

CD4 T cells display remarkable plasticity, responding to pathogen specific cytokine milieus to fine tune responses to mediate protection^44^. It is becoming increasingly recognised that responding fates are not fixed, and in malaria, CD4 T cell fates have been shown to change between subsets and have hybrid states ^45,46^. We have shown in controlled human malaria infection that Tfh/Th1 and Tr1 cells overlap clonally and phenotypically ^10^, while in areas of intense malaria transmission in children, cell phenotypes appear less plastic and skew strongly to Tr1 phenotypes which dominate the response ^7,8^. Additionally, there is significant diversity within the Tfh cell compartment, with specific subsets responding to infection in context dependent manner ^19^. Here we expand on our understanding of CD4 and Tfh cell diversity, showing that Tr1 and Tfh cells within the CD4 T cell compartment, and Tfh10 cells within the Tfh cell compartment, dominate the response in malaria. Identified overlaps between Tr1 and Tfh are consistent with findings from pregnant women, where malaria specific Tr1 cells can also co-produce the Tfh associated cytokine IL-21, and that these cells expand during malaria in pregnancy ^29^. It remains unknown whether the overlapping Tr1 and Tfh cell fates are due to cells with Tr1 fates taking on Tfh markers, or by Tfh cells adopting regulatory programs. However, Tfh10 signatures have been identified within bonafide Tfh cells within human germinal centres in other settings ^47^ ^25^. Further, in animal models of chronic infection with LCMV, Tr1 cells can develop from antigen-specific Tfh cells ^27^, and Tr1 cells emerge from Tfh cells in the context of repeated antigen stimulation ^26,30,48–50^. These development pathways involve epigenetic modifications and changes in the expression of Tfh stabilising and Tr1-promoting transcription factors ^26^. It is possible that the abundance of Tr1 cells within the periphery in malaria is due to ‘spill over’ of regulatory phenotypes of Tfh within germinal centres within secondary lymphoid tissues such as the spleen. Alternatively, Tr1 cells within the periphery may upregulate CXCR5 and seed responses within the germinal centres. Due to low numbers of clonally identified cells in our current study, we are not able to explore development pathways between subsets, and further studies leverage secondary lymphoid tissue cells are required to establish whether Tr1 regulatory signatures can be found within germinal centre Tfh cells in human malaria.

For Tr1 cells, a key role in malaria has been identified, with recent studies in a highly exposed population of Ugandan children showing that these cells have a role in tolerogenic immunity, controlling immunopathology and preventing symptoms at the expense of parasite clearance ^8^. However, how Tr1 like cells may function within the Tfh cell compartment is unknown. In animal models, Tfh derived IL-10 is required to sustain germinal centre reactions in chronic viral infection ^46^ and in mice malaria models, the deletion of IL10R on B cells lead to reduced GC B cell differential and antibody production suggesting that intrinsic B cell IL-10 is essential in malaria humoral immunity ^49^. IL-10 also drives switching of B cells to IgG3 ^52^, a key cytophilic cytokine for protection from malaria. However, given that BLIMP1, required for Tr1 conversion, antagonises Bcl6, required for Tfh cells, how these cells co-exist phenotypically is unclear, however may indicate that the Tr1-like signature dominating in malaria may restrict robust Tfh cells function within the germinal centre. Indeed, antibody responses in malaria develop slowly, only reaching protective levels in adults after multiple infections, consistent with a negative role of Tfh10 within humoral immunity. For both Tr1 and Tfh10 cells, maintenance was associated with repeated antigen exposure, with frequencies of malaria-specific cells wanning in the absences of continuous exposure. Waning was not associated with age, unlike Th1 and Tfh1 cells, which declined more slowly in adults, and more rapidly in small children. This finding is strongly suggestive that regulatory T cells require recent or constant antigen exposure for maintenance, consistent with previous studies that showed associations between the frequencies of Tr1 cells and recent malaria ^8,12^. Additionally, this age-independent waning of Tr1 and Tfh10 are in contrast to the age dependent wanning of humoral immunity reported previously in this same cohort ^37^. Together these data suggest that Tr1 and Tfh10 are a response to antigen, but may not have direct roles in humoral immunity. Further mechanistic studies, including investigating cells directly within secondary lymph nodes during infection are required.

### Limitations

Studies used here identified antigen-specific cells using AIM assays. While this approach allows the study malaria-specific CD4 T cells without relying on tetramers, which face challenges in malaria due to *P. falciparum* antigen diversity ^51^ and the HLA class II gene polymorphisms ^52,53^, non-specific cytokine activation of cells within stimulations are possible, and not all AIM+ cells maybe truly antigen specific ^36^. Additionally, while we used LAG3/CD49b to identify Tr1 cells based on previous studies, recent work has shown that Tr1 malaria specific cells can be more definitively identified as CXCR6+CD127low ^7,8^. These differences in approaches may underpin differences between findings here and previously in similar study populations ^7,8,13,14,41^. Further, while we identify Tr1/Tfh cell overlaps here, bonafide Tfh cells can only be found with secondary lymphoid tissues ^18,54^. While clonal overlap between Tfh cells with the secondary lymphoid compartment and the blood have been identified ^55,56^, further studies in these secondary lymphoid compartments are required to dissect whether Tfh10 are functional with the germinal center in malaria.

## Materials and methods

### Ethics statement

Written informed consent was obtained from the parent or guardian of all the children and from adult study participants. The study protocols were approved by the Uganda National Council of Science and Technology (HS 1019), Makerere University School of Medicine Research and Ethics Committee (2011-167), University of California, San Francisco Committee on Human Research (11-05995), QIMR Berghofer Human Ethics Committee (P3444) and Alfred Hospital Human Ethics Committee (188/23).

### Study site and population

Samples were collected from a longitudinal cohort of children and adults in the East African International Centres of Excellence in Malaria Research (ICEMR) study in Nagongera sub-county, Tororo district, Uganda ^31^. Tororo had one of the highest malaria burdens of all surveyed study sites nationally, having year-round transmission with peaks occurring in October to January and April to July ^57^. Briefly, the ICEMR cohort enrolled 100 randomly selected households (PRISM1), with all children enrolled in the study if they met the following criteria, (i) aged above 0.5 and less than 10 years (ii) full time resident in the household, (iii) no intention to move out of the sub county in the next 2 years, (iv) agreement to come to the study clinic for any febrile illness, (v) agreement to avoid antimalaria medication administered outside the study (vi) provision of written informed consent from a parent or guardian. One adult caregiver capable of providing informed consent was also enrolled ^31^. Study participants presenting with fever (tympanic temperature >38 °C) or a history of fever in the last 24 hours, were tested by blood smear for *Plasmodium* parasites. If found positive, they were treated with artemether-lumefantrine, regardless of the parasite density, as per the national guidelines. Participants were routinely assessed for malaria parasites at the clinic every three months and additional blood was collected for immunology studies. All malaria negative samples by microscopy were further tested for sub microscopic malaria parasites using a molecular test, loop-mediated isothermal amplification (LAMP) ^31^. Prior to December 2014, malaria control measures were limited to the distribution of long-lasting insecticide-treated bed nets and case management with Artemisinin-Based Combination Therapy. In December 2014, indoor-residual spraying (IRS) with the carbamate bendiocarb was initiated, with additional rounds administered in June 2015 and November 2015. In June 2016, IRS was administered with the organophosphate pirimiphos-methyl (Actellic), with repeated rounds in June 2017 and June 2018. IRS resulted in dramatic reduction in malaria transmission ^58^.

PBMCs were collected from blood drawn at routine visits via density centrifugation with Ficoll-Paque before cryopreservation. For the current study, PBMCs were were selected based on sample availability. For each child participant, individuals were chosen based on availability of 4 repeated samples over 8-18 month period where individuals were asymptomatically infected, but with at least one symptomatic malaria episodes between blood draws. scRNAseq samples for adults were selected from another sub study in the same study site (PRISM2) within the ICEMR investigating malaria dynamics after a period of IRS ^58^.

### *Plasmodium falciparum* culture

*P. falciparum* 3D7 parasites were cultured *in vitro* in human O+ red blood cells at 5% haematocrit in RPMI media containing 5% heat-inactivated human sera, 0.25% AlbuMAX II (Gibco), 30 ug/mL gentamicin (Gibco), 25 mM HEPES (Gibco) and 370 µM hypoxanthine (Sigma). Parasites were grown at 37 °C in a gas mixture of 94% N_2_, 5% CO_2_, 1% O_2_ and maintained to the mid-late trophozoite stage. Trophozoites were enriched using Magnetic Activated Cell Sorting columns MACS® CS and the purified parasite-infected red blood cells were cryopreserved in Glycerolyte 57 Solution (Baxter).

### Single cell RNA sequencing of CD4 T cells and Tfh cells

#### Parasite stimulation and cell sorting

To enrich for malaria specific cells, PBMCs were thawed in RPMI 1640 media supplemented with L-Glutamine, 25 mM HEPES,10% Fetal calf serum (FCS) and 0.02% benzonase nuclease. Briefly, 3-4 x10^6^ PBMCs per donor were added to 96-well U- bottom plates at a concentration of 1 x10^6^ cells / 200 µL media per well. Cells were rested for 2 hours at 37°C in 5% CO_2._ After rest, the cells were stimulated with 3D7 *P. falciparum* infected red blood cells (pRBC) at a 1:1 ratio of PBMC: pRBC for 18 h. We enriched CD4^+^T cells using a negative selection CD4 T cell isolation kit (Miltenyi Biotec) through a QuadroMACS*^TM^* Separator (Miltenyi Biotec; Cat. #130-090-976), with pre-Separation 158 Filters (Miltenyi Biotec; Cat. #130-041-407) and LS Columns (Miltenyi Biotec; Cat. #130-042-401) according to the manufacturer’s instructions. The CD4 T cells were stained at room temperature with the following antibodies; anti-CD4 PerCP-Cy5.5 (OKT4), anti-CD45RA BB515 (H100), anti-CD69PE (FV50), anti-OX40 APC (ACT35) and viability dye SYTOX^TM^ Blue (Invitrogen). The stained CD4 T cells were sorted using the FACSAria III (BD Biosciences). Populations were gated as live singlets CD4^+^CD45RA^-^ and sorted as either OX40^+^CD69^+^ (AIM+) or OX40^-^CD69^-^ (AIM-) (Supplementary Fig S1A, Supplementary Table S6).

For Tfh cells, PBMCs were stimulated as for CD4 T cells and stained with anti-CD4 PerCP-Cy5.5 (OKT4), anti-CD45RA BB515(H100), anti-CXCR5 BV711 (J252D4) and viability dye SYTOX^TM^ blue (Invitrogen). Tfh cells were sorted as live CD3^+^CD4^+^CD45RA^-^CXCR5^+^ into 2% FCS/PBS on BD FACSAria III Cell sorter (Supplementary Table S6, Supplementary Figure 1A). After sorting, all AIM⁺ and approximately 6000 AIM⁻ CD4 T cells per donor were combined. The CD4T cells and Tfh cells were then pooled by time point, with each time point loaded onto a separate lane of the 10x Chromium Next GEM chip. Cells from adults were loaded across timepoints.

#### Genomics Chromium GEX /VDJ Library preparation and sequencing

Gel Bead-in-Emulsion (GEMs) were generated in Chromium Controller and processed using the Chromium Next GEM Single cell 5’ GEM Kit v2 PN 1000244 (Chromium Next GEM Chip K Single Cell Kit PN 1000286, Gel Bead Kitv2 PN 1000264, Library Construction Kit PN 1000190, TCR Amplification Kit PN 1000252, Dual Index Kit TT Set A PN 1000215). Samples were run in two batches and data integrated for subsequent analysis. Expression libraries were generated as per the manufacturer’s instructions. Library quality and concentration were assessed using the Agilent TapeStation with High Sensitivity D5000 Screen Tape Assay and Qubit Fluorometer with dsDNA BR assay kit according to the manufacturer’s instructions.

The 5’ Gene Expression library and V(D)J libraries were pooled at 4:1 ratio respectively and sequenced using Nextseq 550 (Illumina) with the following parameters; paired-end dual index sequencing (150 bp Read 1 and 150 bp Read 2), Read 1:26 cycles, i7 index:10 cycles, i5 index: 10 cycles and Read 2: 90 cycles. A minimum of 37,000 reads pairs per cell was targeted.

#### Demultiplexing and analysis

The Illumina BCL files were processed using Cell Ranger version 3.1.0 software (10x Genomics). The FASTQ files reads were aligned to the GRCh38 human genome reference using the default parameters. Cell Ranger count matrices for each sample (donor and timepoint) were merged and analysed using Seurat package v4.3.0. Data from the pooled donors was demultiplexed using Variational Inference for Reconstructing Ensemble Origin (VireoSNP)^59^. Donor identities were predicted using the cell’s haplotype ratios, and only cells identified as singlets were included in the downstream analysis ^9,59^. For quality control, cells with less than 50 genes (low quality) or greater than 6,000 genes (doublets) and greater than 9% mitochondrial genes were removed. Reads mapping to the TBR and RPL genes were also removed. The filtered data sets were normalised using the SCTransform function ^60^ and effects of mitochondria DNA transcripts were regressed out by the *vars.to. regress* parameter. Variations due to batch, were corrected using the integration workflow in Seurat v4.3.0. and the data split into the CD4 T cell and Tfh data sets.

For CD4 T cells malaria-specific cells were identified based on the expression of *CD69* and *TNFRSF4* (which encodes OX40) and classified as AIM+(*CD69*>0 and *TNFRSF4* >0) or AIM-. The dataset was split by age group (Child/Adult) and AIM status. Subsequently, each agegroup_AIM dataset was normalised using SCTransform ^60^. The shared highly variable genes across datasets were determined using the FindIntegrationAnchors function and then used to integrate the data sets with the IntegrateData function. Principal component analysis (PCA) was carried out using RunPCA, and the first 20 PCs were used to generate UMAP visualisation using RunUMAP. For Tfh cells, the data set was split by donor and each donor data set was normalised using SCTransform ^60^. The shared highly variable genes across datasets were determined using the FindIntegrationAnchors function and then used to integrate the data sets with the IntegrateData function. Principal component analysis (PCA) was carried out using RunPCA and the first 20 PCs were used to generate UMAP visualisation using RunUMAP.

#### Cell clustering and annotation

The top 30 principal components (PCs) were calculated using the RunPCA function and used as the number of dimensions for the FindNeighbors and FindClusters functions. To find the optimal clustering resolution, we explored a range of resolutions from 0.1-1.4 and 0.2- 0.5 for CD4 T cells and Tfh cells respectively. The range of clusters was visualised on a cluster tree using the R package clustree (v0.43). Resolutions 1.2 with 16 clusters and 0.5 with 8 clusters were selected for downstream analysis of CD4 T cells and Tfh cells respectively. Clusters were visualised in two-dimensional space using the Uniform Manifold Approximation and Projection (UMAP). Cell clusters were annotated based on canonical lineage and functional genes generated by ‘FindAllMarkers’ function in Seurat. Malaria specific Tfh cells were identified by expression of CD69 with either *TNFRSF4 or, IL2RA or TNFRSF9* (encoding OX40, CD25 and CD137 respectively).

Due to a wetting error failure on the 10X Chromium, the cell recovery of the Timepoint (T0) for CD4 and Tfh data sets was very low and while these cells are included in the clustering, they were excluded from the proportion analysis.

#### TCRA/B clonal analysis

Paired chain TCR sequences were obtained by targeted amplification of full-length V(D)J segments during the library preparation. The sequence assembly and clonotype calling were performed using Cell Ranger version 3.1.0 software (10x Genomics) with the default human V(D)J reference database. Filtered TCR annotations were integrated with matched single-cell gene expression data using the R package scRepertoire version 2.0.0 ^61^, which assigns TCR nucleotide and amino acid sequences, clonal frequency counts and clonotype classification to each cell. Only cells with complete paired TCRα and TCRβ chains were included in the clonotype analysis. Unique clonotypes were identified based on shared identical amino acid sequences in the complementarity-determining region (CDR3) of the TCRα and TCRβ chains. TCR clonal diversity was assessed using Simpson’s diversity index, with diversity reported as 1 − D, where D is the sum of squared clonal proportions.

### Activation Induced Marker (AIM) assay for flow cytometry

To assess antigen specific CD4 T cells and Tfh cells at protein level, we performed AIM assay with parasite infected RBCs and CMV peptide pools as previously described ^33^. PBMCs were thawed in RPMI 1640 media supplemented with L-Glutamine, 25 mM HEPES,10% Fetal calf serum (FCS) and 0.02% benzonase nuclease. The cells were resuspended at a concentration of 1 x10^6^ cell/ 200µL media per well in 96 well plates and rested for 2 h at 37 °C in 5% CO_2._ After rest, the cells were stimulated with either parasite infected red blood cells (pRBC) or uninfected red blood cells (uRBC) at a 1:1 ratio of PBMC: pRBC/uRBC or 0.5 mg/ml of overlapping peptide pools for CMV pp65 (Cat# PM-PP65) for 18 h. Unstimulated cells were included as negative controls. After stimulation, the cells were incubated for 45 min with human Fc Block (BD Biosciences) and anti-CCR7, anti-LAG3, and anti-CD49b antibodies. The cells were washed with PBS and surface stained at room temperature with Live/Dead BLUE viability stain (Invitrogen) and antibodies from BD Biosciences and BioLegend (Supplementary Table S7). After washing with 2% FCS/PBS, the cells were fixed with BD stabilising solution for 15 min at room temperature. The fixed cells were then resuspended in 2% FCS/PBS, and data were acquired using a 5-laser Cytek Aurora spectral flow cytometer

### Flow cytometry data analysis

Flow cytometry data was analysed using Flowjo V10.7.1 (BD Biosciences) and R v4.1.3 using the Spectre R package (v1.0) ^62^. Antigen specific and non-specific nnCD4 T cells were gated in Flowjo V10.7.1 and a Boolean “NOT” gate used to exclude Tr1 cells based on LAG3 and CD49b expression (Supplementary Figure 4). Data was imported into Spectre as raw channel value CSV files and merged. Batch-derived technical variations in the data were corrected using COARSE alignment function using the 95th percentile of reference samples and applying this adjustment to the samples in each batch. The reference samples were obtained from malaria naïve Australian adults. CD4 T cell subsets were identified by supervised clustering using the Flow Self-Organizing Map (FlowSOM) algorithm ^63^ of 6,251,885 cells. For FlowSOM, random seed was set as default while metaclusters k-value set at 60 to enable over-clustering via the R Spectre (v1.1.0)^62^. Cells were clustered based on expression of CXCR5, CXCR3, CCR6 CCR4, CRTh2, CCR10 and CD45RA. Each cell was assigned to a specific cluster and metacluster. For visualisation 50000 cells per stimulation condition were projected on dimensional reduction using UMAP ^64^.. The summary data generated for all cells was exported to R for further analysis

### Tweedie Mixed Effect Model

Tweedie mixed effects models have been applied in various fields to analyse skewed and complex longitudinal data structures. These models can accommodate non-negative continuous data with a discrete probability mass at zero, making them suitable our data ^65^. The model residuals were assessed using the DHARMa residual test with no significant violations of model assumptions detected ^66^. The subject-level random effects of the model were normally distributed suggesting that changes in the cell frequency were not donor driven.

The Tweedie mixed effect model used to assess the effect of age (in years, centered at 10 years), time (in weeks), and their interaction on the frequency of malaria-specific cells can be described as follows:

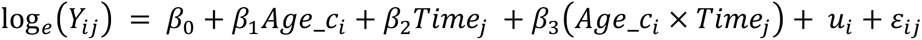

- log*_e_*(*Y_ij_*) – the natural log of the outcome for subject *i* at time *j*
- *β_0_* – intercept

- The expected log(outcome) at *Time_j_* = 0 and *Age*_*c_i_* = 0
- *β*_1_*Age*_*c_i_* – fixed effect of age

- Represents the average change in outcome per one-year increase in age, holding time constant at zero weeks (*Time_j_* = 0)
- *β*_2_*Time_j_* – fixed effect of time

- Represents the average change in outcome per one-week increase in time, holding age constant at the age of 10years (*Age*_*c_i_* = 0)
- *β*_3_(*Age*_*c_i_* × *Time_j_*) – fixed interaction effect

- Represents the average change in the effect of time on the outcome for each one-year increase in age
- 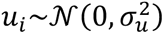 – the random effect for subject *i* (assumed to follow a normal distribution with a mean of zero and variance of 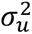)

- Represents subject-level variation from the population-level effects
- ε*_ij_* – residual error term (assumed to follow a compound-Poisson-Tweedie distribution)

To visualize model outputs the glmmTMB package(v 1.1.7) ^67^ was used to fit models with the visreg package(v 2.7.0) ^68^ used to visualise the estimated relationship between time and frequency of malaria-specific cells, conditional on select ages.

### Statistical Analysis

Statistical analyses were conducted using R v4.1.3. Background-subtracted net values were used to assess antigen-specific CD4+ T cells as a percentage of total nnCD4+ T cells. The data was adjusted by subtracting the responses to uRBC from pRBC, or media stimulations from CMV. Comparisons between children and adults were tested using the Wilcox test. Comparisons between malaria and CMV in each individual were tested using paired Wilcox test.

## Data and code availability

For scRNA/TCRseq, cell ranger expression and vdj output is available here: https://www.ncbi.nlm.nih.gov/geo/query/acc.cgi?&acc=GSE279685

Code to reproduce the analysis can be found here: https://github.com/Boyle-Lab-CRDV/tfh-tr1-overlaps-in-malaria.git.

Processed scRNA/TCRseq data is available here: https://doi.org/10.5281/zenodo.20619125

## Author contributions

MN, DA, MS, JRL, ND generated data, supervised by CE and MJB

MN, DAO, ZP analysed data, supervised by CE and MJB

KM, NF, IS, JR, EA, MK, MF, PJ drove clinical cohorts and contributed key resources

MJB, MF and PJ conceptualised the study.

MJB, MF, PJ contributed funding,

MN and MJB drafted manuscript with feedback from all authors.

## Declarations of interests

All authors declare no conflicts of interest

## Acknowledgements

Burnet Institute, University of Melbourne and QIMR-Berghofer acknowledge the traditional custodians of the lands where they are located, the Boonwurrung and Wurundjeri Woi-wurrong people of the Kulin Nation, and the Turrbal and Jagera people.

RBC and human serum were provided by the Australian Red Cross Blood Bank (Brisbane). We thank Stacey Llewellyn for expertise and advise in Tweedie Model development.

## Funding

The parent PRISM study was funded by National Institutes of Health as part of the International Centers of Excellence in Malaria Research (ICMER) program (U19AI089674). This work was supported by the National Health and Medical Research Council of Australia Career Development Award 1141278, Project Grant 1125656, and Ideas Grant 1181932 to MJB); the CSL Centenary Fellowship and the Snow Medical Foundation Fellowship 2022/SF167 to M.J.B; The Burnet Institute is supported by the NHMRC for Independent Research Institutes Infrastructure Support Scheme and the Victorian State Government Operational Infrastructure Support.

## Notes

### Competing Interest Statement

The authors have declared no competing interest.

