## Supplementary Materials for "Circulating T-follicular helper and type I regulatory T cells have overlapping phenotypes in *P. falciparum* malaria and are maintained by parasite exposure"

**Supplementary Figures and Tables**

**Supplementary Tables**

***Supplementary Table 1****:* Cluster markers for CD4 T cell subsets *attached as excel*

***Supplementary Table 2:*** DEGs between Tr1 and Tfh cell subsets *attached as excel*

***Supplementary Table 3:*** Cluster markers for Tfh cell subsets *attached as excel*

***Supplementary Table 4:*** Tweedie model outputs for longitudinal analysis of CD4 T cell frequencies *attached as excel*

***Supplementary Table 5:*** Tweedie model outputs for longitudinal analysis of Tfh cell frequencies *attached as excel*

***Supplementary Table S6:*** Antibodies for parasite specific CD4 T cells and Tfh cell sorting

| **Fluorophore** | **Marker** | **Clone** | **Catalogue number** | **Company** | **Dilution** |
|---|---|---|---|---|---|
| Parasite specific and non parasite CD4T cell sort | | | | | |
| BB515 | CD45RA | HI100 | 564552 | BD | 1/400 |
| AF700 | CD3 | SK7 | 344822 | Biolegend | 1/50 |
| PE | CD69 | FN50 | 310904 | Biolegend | 1/50 |
| PerCPCy5.5 | CD4 | OKT4 | 317428 | Biolegend | 1/50 |
| APC | OX40 | ACT35 | 563473 | BD | 1/50 |
| Tfh cell sort | | | | | |
| BB515 | CD45RA | HI100 | 564552 | BD | 1/400 |
| AF700 | CD3 | SK7 | 344822 | Biolegend | 1/50 |
| PerCPCy5.5 | CD4 | OKT4 | 317428 | Biolegend | 1/50 |
| BV711 | CXCR5 | J252D4 | 563922 | Biolegend | 1/50 |

**Supplementary Table S7:** ***Antibodies for the AIM assay cell phenotyping***

| **Fluorophore** | **Marker** | **Clone** | **Catalogue** | **Manufacturer** | **Dilution** |
|---|---|---|---|---|---|
| Surface staining at RT | | | | | |
| BUV395 | CCR10 | IB5 | 565322 | BD | 1/100 |
| BUV496 | CD8 | RPA-TS | 612942 | BD | 1/100 |
| BUV563 | CD45RA | H100 | 565702 | BD | 1/400 |
| BUV737 | CD69 | FN50 | 612817 | BD | 1/100 |
| BUV805 | CD3 | SK7 | 612893 | BD | 1/50 |
| BV510 | Dump (CD14/ CD19) | MSE2/H1B19 | 30184 | Biolegend | 1/100 |
| BV605 | CCR4 | L291HL1 | 301842 | Biolegend | 1/50 |
| BV650 | CCR6 | 11A9 | 359418 | Biolegend | 1/50 |
| BV711 | CXCR5 | J252D4 | 563922 | Biolegend | 1/50 |
| BV785 | CD4 | RPA-T4 | 300554 | Biolegend | 1/50 |
| PE | CD25 | BC96 | 317442 | BD | 1/200 |
| PE-CF594 | CXCR3 | IC6/CXCR3 | 302906 | Biolegend | 1/50 |
| PE-Cy7 | PD-1 | EH12.1 | 562451 | BD | 1/50 |
| APC | OX40 | ACT35 | 561272 | BD | 1/50 |
| AF700 | CD161 | HP-3G10 | 563473 | Biolegend | 1/50 |
| APC-Fire | CD294/ CRTh2 | BM16 | 350134 | Biolegend | 1/25 |
| LIVE/DEAD Blue | | | L23105 | Invitrogen | 1/5000 |
| staining at 37C | | | | | |
| BV421 | LAG-3 | T47-530 | 565721 | BD | 1/50 |
| FITC | CD49b | HMa2 | 11-0491-82 | eBiosciences | 1/100 |
| PerCP-Cy5.5 | CCR7 | G043H7 | 353220 | BD | 1/25 |

### Supplementary Figures

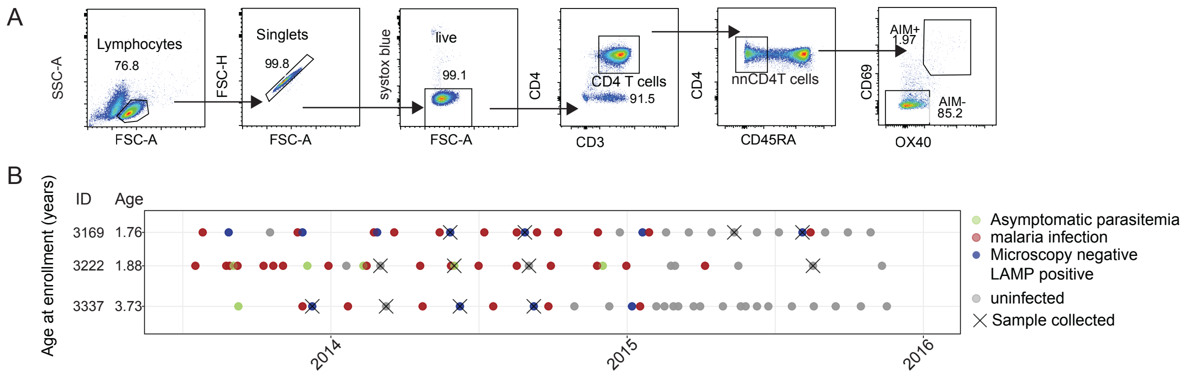


***Supplementary Figure S1: Identification of CD4 T cells and malaria incidence in children.*** ***(A)*** *Flow cytometry gating strategy for identifying AIM$^{+}$ and AIM$^{-}$ CD4$^{+}$ T cells for scRNA/TCRseq.* ***(B)*** *Timelines for malaria incidence and asymptomatic infection in children used in analysis. Each dot represents a clinic visit coloured by malaria status, and "X" indicates the timepoints at which the samples used in this study were collected.*

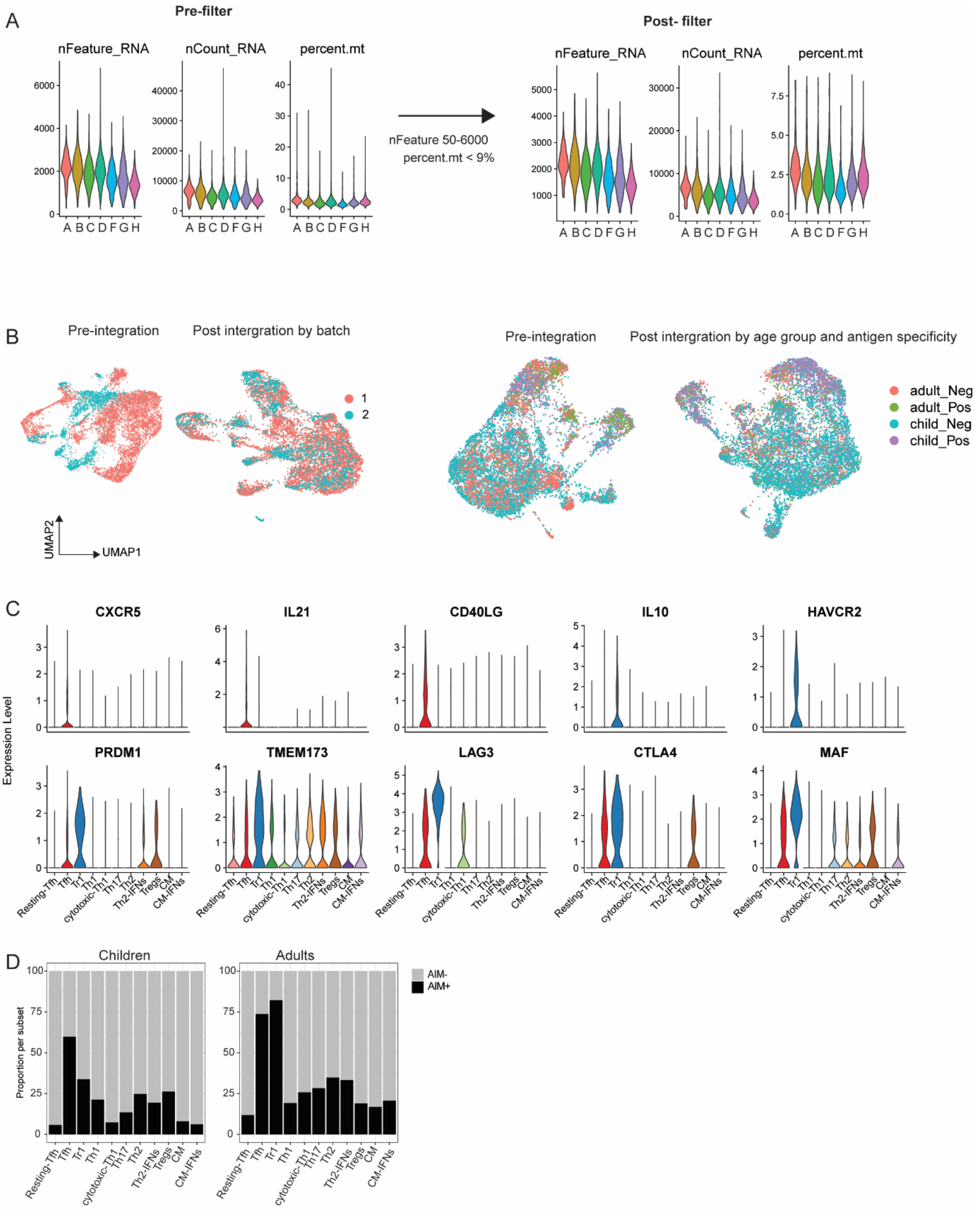


***Supplementary Figure S2: Single cell RNA sequencing of CD4 T cells.***
***(A)*** *Quality control with the pre-filter and post-filter violin plots of genes detected per cell(nFeature_RNA), total number of molecules (UMIs) detected within a cell (nCount_RNA) and reads from mitochondrial genes (percent.mt). Cells were filtered based on nFeature_RNA >50 and < 6000 and percent.mt >9%.* ***(B)*** *Integration for batch effects pre and post integration UMAPs.* ***C)*** *Violin plots showing the expression of Tfh and Tr1 related genes across all cell subsets.* ***D)*** *Proportion of AIM+ parasite specific cells within each cell cluster by age group.*

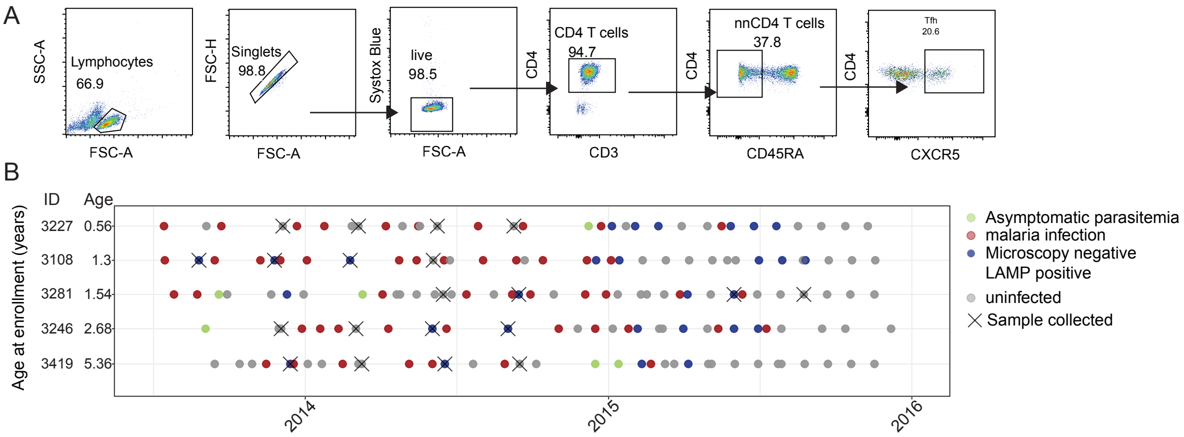


***Supplementary Figure S3: Identification of Tfh cells and malaria incidence in children.***
***(A)*** *Flow cytometry gating strategy for identifying Tfh cells for scRNA/TCRseq.* ***(B)*** *Timelines for malaria incidence and asymptomatic infection in children used in analysis. Each dot represents a clinic visit coloured by malaria status, and "X" indicates the timepoints at which the samples used in this study were collected.*

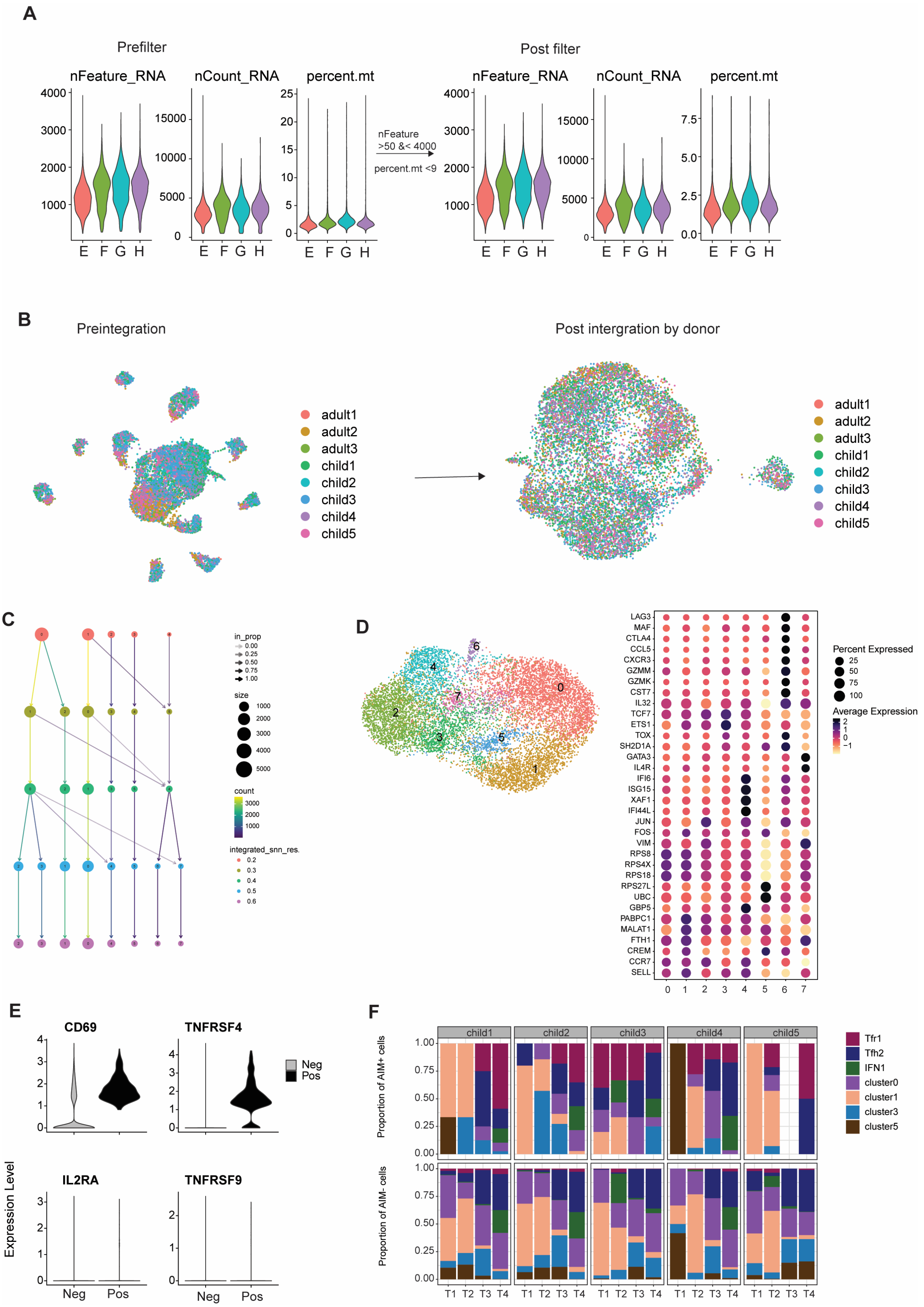
A
Prefilter
nFeature_RNA
nCount_RNA
percent.mt
nFeature
>50 &< 4000
percent.mt <9
Post filter
E F G H
B
Preintegration
Post intergration by donor
adult1
adult2
adult3
child1
child2
child3
child4
child5
C
in_prop
size
count
integrated_snn_res.
D
LAG3
MAF
CTLA4
CCL5
CXCR3
GZMM
GZMK
CST7
IL32
TCF7
ETS1
TOX
SH2D1A
GATA3
IL4R
IFI6
ISG15
XAF1
IFI44L
JUN
FOS
VIM
RPS8
RPS4X
RPS18
RPS27L
UBC
GBP5
PABPC1
MALAT1
FTH1
CREM
CCR7
SELL
Percent Expressed
Average Expression
E
CD69
TNFRSF4
IL2RA
TNFRSF9
Neg
Pos
Expression Level
F
child1
child2
child3
child4
child5
Proportion of AIM+ cells
Proportion of AIM- cells
T1 T2 T3 T4
Tfr1
Tfh2
IFN1
cluster0
cluster1
cluster3
cluster5

***Supplementary Figure S4: scRNAseq of Tfh cells***

***(A)*** *Quality control with the pre-filter and post-filter of genes detected per cell(nFeature_RNA), total number of molecules (UMIs) detected within a cell (nCount_RNA) and reads from mitochondrial genes (percent.mt). Cells were filtered based on nFeature_RNA >50 and < 4000 and percent.mt >9%.* ***(B)*** *UMAP visualisation before and after integration by donor.* ***(C)*** *Cluster tree used to select the optimal resolution for clustering.* ***(D)*** *UMAP of integrated cells clustered using Seurat's FindClusters function at a resolution of 0.5, yielding eight clusters. Clusters 2 and 7 were combined during annotation. Dot dot plot shows expression of canonical and functional genes used to annotate each cluster.* ***(E)*** *Violin plots showing the expression of the genes used to classify malaria specific cells. Malaria specific specific cells were identified based on expression of CD69 >0 with either TNFRSF4 >0 or IL2RA >0 or TNFRSF9.* ***(F)*** *Proportion of malaria specific (AIM+ and non-malaria specific (AIM-) Tfh cell subsets per child across the four time points*

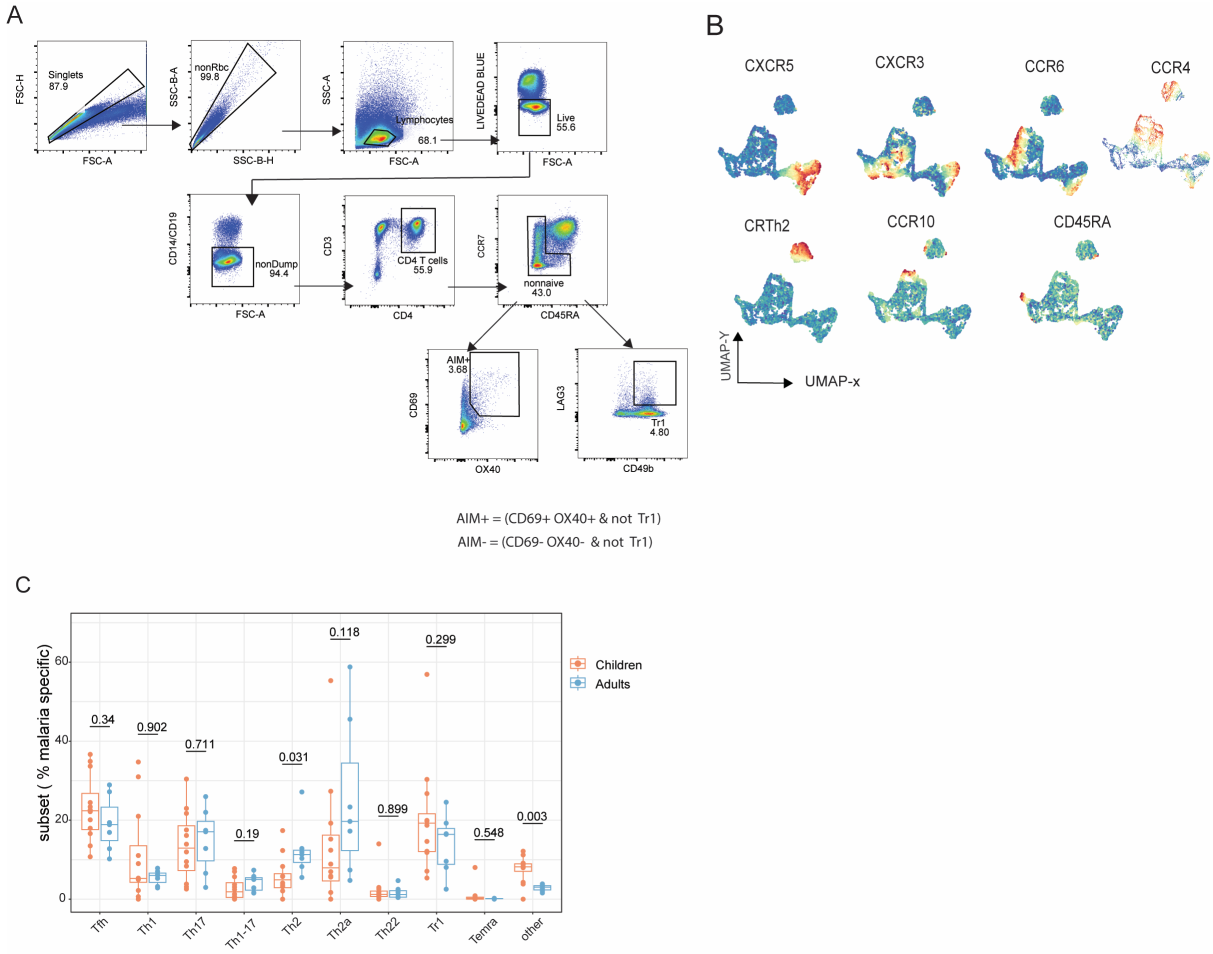


***Supplementary Figure 5: Flow cytometry analysis of antigen specific CD4 T cells in malaria exposed children and adults**. PBMCs from children (n=12) and adults (n=7) stimulated with either Pf infected or uninfected red blood cells and CMV peptides or media, stained and analysed by flow cytometry. **(A)** Flow cytometry gating strategy to identify AIM+ and AIM- CD4 T cells based on co-expression of CD69 and OX40 exported for analysis using SpectreR package. Tr1 cells were manually gated and Boolean "AND/NOT" gates used to select antigen specific (AIM+) nnCD4T cells excluding Tr1 cells from Spectre R analysis. **(B)** Relative expression of the markers used to identify CD4 T cell subsets with the red representing areas of high expression. **(C)** Malaria specific CD4 T cell subsets as proportions of nnCD4 T cells cultured with no stimulation. Box plots show the median and IQR of donors, group comparisons performed by Wilcoxon test with p>0.05 significant.*

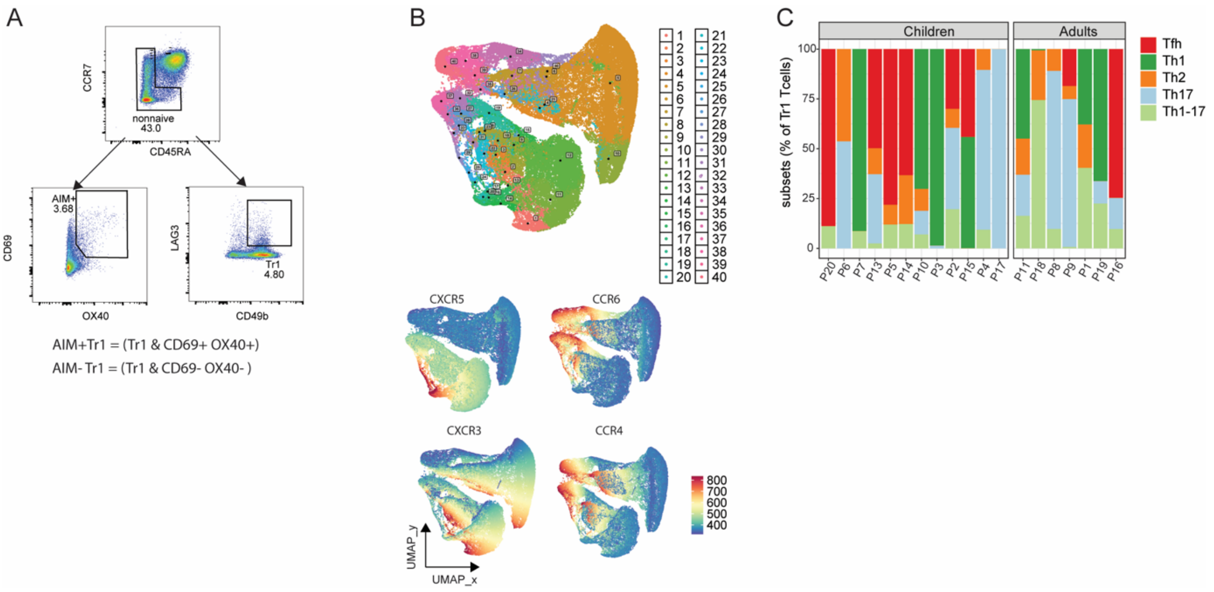


***Supplementary Figure 6: Flow cytometry analysis of Tr1 cells in malaria-exposed children and adults.***

***A)*** *Flow plots showing the manual gating of Tr1 cells.* ***(B)*** *UMAP of Tr1 cells and expression of markers used in clustering.* ***(C)*** *Composition of Tr1 cells across donors.*
